# Alleviating cell-free DNA sequencing biases with optimal transport

**DOI:** 10.1101/2024.04.04.588204

**Authors:** Antoine Passemiers, Tatjana Jatsenko, Adriaan Vanderstichele, Pieter Busschaert, An Coosemans, Dirk Timmerman, Diether Lambrechts, Daniele Raimondi, Joris Robert Vermeesch, Yves Moreau

## Abstract

Cell-free DNA (cfDNA) is a rich source of biomarkers for various (patho)physiological conditions. Recent developments have used Machine Learning on large cfDNA data sets to enhance the detection of cancers and immunological diseases. Preanalytical variables, such as the library preparation protocol or sequencing platform, are major confounders that influence such data sets and lead to domain shifts (i.e., shifts in data distribution as those confounders vary across time or space). Here, we present a domain adaptation method that builds on the concept of optimal transport, and explicitly corrects for the effect of such preanalytical variables. Our approach can be used to merge cohorts representative of the same population but separated by technical biases. Moreover, we also demonstrate that it improves cancer detection via Machine Learning by alleviating the sources of variation that are not of biological origin. Our method also improves over the widely used GC-content bias correction, both in terms of bias removal and cancer signal isolation. These results open perspectives for the downstream analysis of larger data sets through the integration of cohorts produced by different sequencing pipelines or collected in different centers. Notably, the approach is rather general with the potential for application to many other genomic data analysis problems.

## 1 Introduction

Cell-free DNA (cfDNA) has been identified as a promising source of biomarkers for the detection of fetal aneuploidy [1, 2], transplant rejection [3], incipient tumours [4], autoimmune disease [5] or inflammatory disease [6]. While cfDNA fragments in healthy individuals primarily originate from the apoptotic release of DNA from cells of hematopoietic origin [7], these fragments can also be of tumoural origin in cancer patients. While most clinical applications of cfDNA in oncology focus on finding tumour mutations (e.g., using a targeted panel of cancer driver variants) [8, 9], a lot of research has been carried out around the analysis of coverage and fragmentome profiles. Indeed, the copy number aberrations (CNAs) carried by the genome of cancerous cells are detectable by low-coverage whole-genome sequencing and downstream analysis of cfDNA from cancer patients [10, 11]. Because cfDNA can be collected in a non- or minimally-invasive manner (e.g., blood draw), and thanks to the cost-effectiveness of shallow whole-genome sequencing, liquid biopsies are a valuable candidate for population-wide cancer screening [7, 4] and diagnosis, and considerable research has been devoted to assessing their clinical utility [12].

Fragmentomic analysis of cfDNA offers the possibility to improve its use as a sensitive biomarker for cancer detection [13, 14], as cfDNA fragments mirror the chromatin accessibility, nucleosome positioning and degradation pattern of their tissue of origin [15, 16]. For this reason, CNA calling can be complemented with fragmentation profile analysis based on fragment length, as well as positional information [13, 17, 18, 19]. Indeed, the distribution of cfDNA fragment lengths is shifted downward in circulating tumour-derived DNA (ctDNA), supporting signatures of their cellular origin [15, 20].

Beyond fragmentomics, methylation patterns are indicative of the tissue of origin, and methylation signatures have been exploited for sensitive cancer detection and tissue-of-origin identification [21]. Finally, recent work has been devoted toward integrating multiple properties of cfDNA within a single multimodal analysis approach, including variant calling, CNAs, methylation and fragmentomic profiles, as well as other complementary sources of information such as nucleosome-depleted region (NDR) profiles [22] or fusion gene detection [23].

However, the development of reliable models that are predictive of relevant clinical outcomes (for example, diagnosis) remains challenging because of the limited number of available cases (especially for disorders with smaller incidence rate), the high dimensionality of cfDNA data and the various sources of biases related to preanalytical settings. These latter biases mainly arise when protocol changes are introduced over time or between different centers. For example, the choice of blood collection tube might affect cfDNA concentrations and the prominence of leukocyte DNA [24, 25], which could in turn affect the detection of low-frequency variations from cancerous cells. Other preanalytical factors include the delay before centrifugation and protocols for plasma separation, and plasma storage conditions [26]. For example, two-step centrifugation reduces contamination by genomic DNA thanks to reduced white blood cell lysis, compared to one-step centrifugation [27]. Moreover, some DNA extraction platforms, such as Maxwell and QIAsymphony, preferentially isolate short fragments over long ones [28]. The choice of library preparation kit directly affects the distribution of read counts, as the polymerase enzymes used in these kits have different levels of efficiency in amplifying fragments with low vs. high GC-content [29]. For instance, some library preparation kits (e.g., Nextera XT) introduce a bias toward low-GC regions [30]. Multiplexed sequencing without suitable dual indexing can result in barcode swapping, and the swapping rates are platform-dependent (e.g., higher on HiSeqX or 4000 compared to MiSeq) [31]. Index swapping mechanism is caused both by multiplex PCR and flow cell chemistry, and is responsible for cross-contamination within the same pool [32]. Finally, the choice of sequencing instrument also plays a role. For example, different GC-content bias profiles have been reported for Illumina MiSeq and NextSeq platforms, compared to PacBio or HiSeq [33].

In this work, we focus on the bias correction of genome-wide copy-number (i.e., GIPseq [4]) profiles based on normalised read counts. The aforementioned preanalytical settings can affect the read counts, for example through differential coverage of regions differing by their GC content, thus invalidating direct statistical analysis (e.g., using *z*-scores) of CNA profiles. Moreover, these distributional shifts [34] are not properly handled by classical Machine Learning algorithms and are responsible for performance drops on test sets. Mitigating these biases is therefore of utmost importance in strengthening biological signals and guaranteeing performance on unseen data. Such a task typically falls in the category of domain adaptation (DA) [35] problems, where computational methods are needed to compensate for the fact that a given model is tested on data drawn from a different distribution than the ones on which it has been trained. In this work, we will refer to the samples being corrected as belonging to the source domain, while the fixed data lies in the target domain. We restricted ourselves to unsupervised DA, where the variable of interest (e.g., whether an individual is affected by a certain condition) is unknown. Such annotations are not necessarily available, especially for rarer diseases. Also, already-existing methods (GC correction) don’t require such information and are widely applicable, as they can be applied in a sample-wise fashion. This is highly relevant due to GC and sequencing biases not only operating at the domain-level, but also at the individual level [36]. When multiple source domains coexist, the problem is referred to as a domain generalisation problem [37]. Since multiple domain shifts occurred in our data sets over time, our own method falls under this category.

Previous work on the bias correction of copy-number profiles has mostly been directed toward GC-content and mappability bias correction. Benjamini and Speed [38] originally categorised these methods as single position models, fragmentation models, read models, full-fragment models and global models. An example of global model is the widely-used LOESS GC-content bias correction [39, 40], which decorrelates the per-bin GC-content percentage from the normalised read counts based on local regressions. `ichorCNA` [41] is a tool for calling CNA from read counts, that internally performs mappability and GC-correction in a similar way. BEADS [36] falls into the category of read models, as it re-weights individual reads based on their GC-content before computing their per-bin counts. The single position model from Benjamini and Speed [38] relies on the computation of the mean fragment count for each GC stratum, by considering all the mappable positions along the genome having similar GC-content. Finally, the LIQUORICE algorithm [23] operates at the fragment-level, by computing a coverage weight for each position covered by each fragment. More recently, distance learning and k-nearest neighbours have been proposed [42] to correct coverage profiles. As opposed to previous work, the latter approach exploits information from the whole data set to correct each individual sample.

On the Machine Learning side, previous work on unsupervised DA includes the following. Discriminator-free domain adversarial learning [43] uses domain adversarial learning [44] to obtain a common representation space for all domains. Kernel mean matching [45] aims at matching the higher-order moments of the underlying distributions using kernel functions. Multilevel domain adaptive learning matches the distributions at each intermediate layer of the neural network in a hierarchical fashion [46]. Reconstruction-based methods, such as Cycle-Consistent Adversarial Domain Adaptation (CyCADA) [47], reconstruct samples from the target domain using the samples from the source domain as input. It should be noted that most existing methods use a latent space to represent the samples, which means that the debiased representation is not directly interpretable, which runs afoul of ubiquitous need for interpretability and explainability in human genetics [48]. A key motivation for our work is thus to design a domain adaptation method that adjusts cfDNA profiles in a transparently interpretable manner, by operating at the read count level (i.e., without having recourse to a latent space as domain adversarial methods would) and preserving the *z*-scores produced by the original data.

In this article, we present an advanced data normalisation method for cell-free DNA sequencing data building on optimal transport (OT) theory [49, 50]. OT builds on strong mathematical bases and allows to define a patient-to-patient relationship across domains without the need to build a common latent representation space, as mostly done in the DA field. This enables high interpretability, as samples can be corrected in the original data space (e.g., read counts) directly. Because we originally designed this approach for the correction of normalised read counts within predefined bins, it falls under the category of “global models” according to the Benjamini/Speed classification [38]. In summary, we aim at correcting and mapping the data distribution from a source domain onto the data distribution obtained in a target domain, to enable more robust downstream analysis. As the ultimate goal is to go beyond the classical case–control setting and build models capable of accurately processing data from various sources, we hypothesised that bias removal is a good candidate to increase the effective size of available data sets through their fusion and thus benefit from the scalability of Machine Learning models and enhance their performance. This flexibility would, among other things, reduce the need for laboratories to consistently build new reference sets, as well as enable high reusability of older samples or data collected in unrelated studies. In this article, we report enhanced cancer detection with prior domain adaptation and show that cohorts can be corrected to match the same distribution while preserving the original biological signals (e.g., copy number aberrations) in each patient.

## 2 Results & discussion

In each of the following experiments, we compared our domain adaptation approach to the original data (no correction), as well as center-and-scale standardisation and LOWESS GC-content bias correction, when relevant. Center-and-scale standardisation consists in standardising the data points from each domain separately, by subtracting their median and dividing by the average squared deviation from the median. As this method is univariate, it has been performed on each bin and each data set separately.

### 2.1 Preanalytical biases can be accurately removed by optimal transport

In Fig. 1A, we performed a (Gaussian) kernel principal component analysis on the controls from the HEMA data set to illustrate the impact of the change in library preparation kit on the coverage profiles.

**Figure 1:**
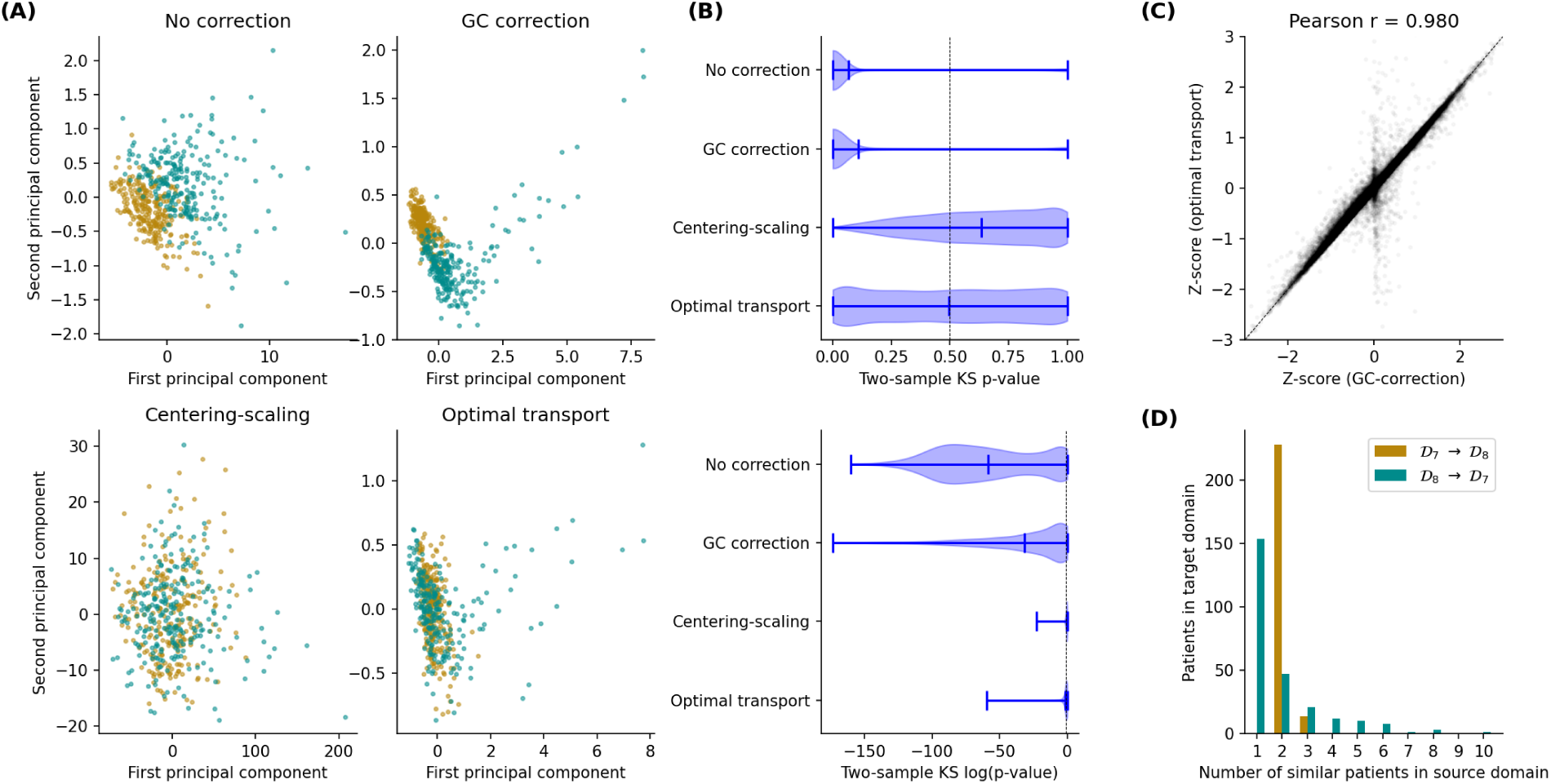
Correction of the healthy samples from the HEMA data set. (A) Kernel principal component analysis of coverage profiles from the two control cohorts (haematological cancer data set). (B) Two-sample Kolmogorov-Smirnov testing, for each bin, of the difference between the two cohorts. *p*-values are shown in both linear and log scales. (C) *z*-scores of the coverage profiles before and after GC-correction and domain adaptation. (D) Histogram depicting, for each patient of the target domain, the number of patients in the source domain for which the transport plan shows a relationship.

The two control sets belong to domains *D*_7_ and *D*_8_ from Table 6, respectively. It appears immediately that LOWESS GC-correction is not sufficient for superimposing the two panels of controls. Center-and-scale standardisation and optimal transport both succeed in that respect, which is expected since these two methods have been designed to explicitly correct data sets. Conversely, GC-correction alleviates GC-content biases at the level of individual samples only.

For each 1 Mb bin, we ran a Two-sample Kolmogorov-Smirnov test (two-sided) to quantify the differences in normalised read counts between the two panels from domains *D*_7_ and *D*_8_, and reported the distribution of per-bin *p*-values in Fig. 1B, both in linear and log scales. While center-and-scale standardisation and optimal transport show similar distributions, the latter contains a median *p*-value close to 0.5 and a more uniform distribution. Such property is desirable since *p*-values are expected to be uniformly distributed under the null hypothesis and some other conditions [51]. Indeed, a distribution shifted leftward indicates the presence of confounders responsible for some discrepancy between the two distributions, as shown both in the absence of correction or with GC-correction. Inversely, a rightward-shifted distribution illustrates overcorrection, as suggested for center-and-scale standardisation. This shift can however also occur for biological reasons, for example when the samples in the first domain are replicates of the samples from the second domain.

In Fig. 1C, we reported the *z*-scores of each sample and bin all together in a single scatter plot, before and after correction. We observe a good consistency between the GC-corrected normalised read counts and the OT-adapted ones, supported by a Pearson correlation coefficient of 0.98. This result suggest that our DA method does not appear to be overcorrecting the normalised read counts. We however observe a very small subset of the bins around 0 for which our method seems to overcorrect the normalised read counts. These bins are located in low-mappability regions, and we suggest that our method is correcting in these few regions relatively more due to the lack of information (the z-scores between these bins and the flanking bins are either not consistent, or mostly made of zeroes). These distortions are irrespective to the original sequencing depth, as the coverage profiles have been normalised. Given the lack of reliability of the original data in these bins (mostly zero counts), we suggest that the information loss is residual compared to high-mappability regions. Let’s note that it is usually advised in the literature to disregard these bins before performing further analysis.

Finally, Fig. 1D reports the number of non-zero entries in the final transport plan Γ inferred by our model, for each patient of domains *D*_7_ (green bars) and *D*_8_ (golden bars). Without the use of entropic regularisation on Wasserstein distance, the model naturally assigns each control from the source domain to *multiple* controls from the target domain, thus reflecting the underlying complexity of the biological processes that generate the read counts. These peculiarities are implicitly acknowledged by our model, by not enforcing the patients to be assigned in pairs.

We conducted similar analyses on the OV and NIPT data sets and obtained slightly different results due to the smaller numbers of samples. In particular, visualisation based on kernel PCA show that the corrected cohorts are still not centered one onto the other. Indeed, since our convergence criterion builds on statistical tests, our algorithm is designed to halt earlier, due to *p*-values being higher when the number of samples is low. This mechanism prevents overcorrection when the available data is insufficient for accurate bias estimation. Also, histograms on the entries of the transport plans showed that each patient from the source was mapped on exactly one patient from the target domain, which met our expectations on these two data sets due to the way domains have been defined (samples were paired). Results have been reported in Suppl. Fig. 2-8.

### 2.2 Patient-to-patient mapping is accurate when the cohorts are representative of the exact same population

While our method is effectively capable of superimposing patients cohorts there is no *a priori* guarantee, besides theoretical considerations, that the coverage profiles are being corrected in the right direction. For this purpose, we considered 64 biological samples with ovarian carcinoma that have been processed with both Illumina HiSeq 2500 and HiSeq 4000 sequencing platforms. The two cohorts belong to domains *D*_10_ and *D*_9_ from Table 6, respectively. In this section, we applied our domain adaptation technique on these two cohorts to see whether the bias removal is decreasing the distance between profiles originating from the same biological sample. Indeed, paired profiles are expected to overlap when the biological variation overcomes technical biases. By design, the cohorts consisted of the same patients, therefore we not only tested our algorithm with default hyper-parameters (noted as “default” in the table), but also without regularisation or early stopping criterion (“*λ* = 0”), and using the transport plan directly to assign pairs and compute accuracy. The purpose was mostly to test the assignment of patients and assess whether OT can map each sample to its correct counterpart.

In Table 1, we compared our correction method with the GC-correction approach, as well as center-and-scale standardisation. Because domain adaptation and estimation of accuracy can be done in two ways (as there are two domains), we reported both settings as separate columns in the table. While center-and-scale standardisation and GC-correction fails at pairing the samples with more than 50% accuracy, we observed a sharp improvement in accuracy with our domain adaptation approach when disabling regularisation. Without correction, only 17 and 14 profiles were correctly assigned to their counterpart in the source domain. GC-correction enabled the correct assignment of 23 and 30 patients, while our domain adaptation approach allowed the correct mapping of 47 and 48 patients when *λ* = 0.

**Table 1:** Performance assessment using paired samples from the OV data set. Accuracy obtained after data correction, when assigning a sample from target domain to the closest sample in source domain, and using the Euclidean metric. In the first column, samples from domain *D*_9_ have been corrected toward *D*_10_, and vice versa.

| Method | Correctly identified pairs |  |
| --- | --- | --- |
| | $\mathcal{D}_9 \rightarrow \mathcal{D}_{10}$ | $\mathcal{D}_{10} \rightarrow \mathcal{D}_9$ |
| No correction | 17 / 64 | 14 / 64 |
| Center-and-scale | 20 / 64 | 27 / 64 |
| GC-correction | 23 / 64 | 30 / 64 |
| Optimal transport (default) | 24 / 64 | 31 / 64 |
| Optimal transport ($\lambda = 0$) | 47 / 64 | 48 / 64 |
| Optimal transport ($\Gamma$) | 51 / 64 | 50 / 64 |

In Table 2, we report the same metric on the NIPT data set, where 563 patients have been sequenced twice with different protocols. This data set has been divided in 6 validation groups and each group divided in 2 domains (see table 6). Each group was designed to control for exactly one preanalytical variable. As an example, the *D*_1*,a*_ and *D*_1*,b*_ domains differ by their library preparation kits, namely

**Table 2:** Performance assessment using paired samples from the NIPT data set. Accuracy obtained after data correction, when assigning a sample from target domain to the closest sample in source domain and using the Euclidean metric, on each of the 6 validation groups. Each validation group was designed to control for one preanalytical variable at a time.

| Preanalytical variable | Comparison | No correction | Center-and-scale | GC-correction | Optimal transport |  |  |
| --- | --- | --- | --- | --- | --- | --- | --- |
| | | | | | $\lambda = 0.5$ | $\lambda = 0$ | $\Gamma$ |
| Lib. prep. kit | TruSeq Nano/Kapa HyperPrep ($\mathcal{D}_{1,a}/\mathcal{D}_{1,b}$) | 2 / 66 | 28 / 66 | 28 / 66 | 44 / 66 | 58 / 66 | 64 / 66 |
| Adapters | Kapa/IDT ($\mathcal{D}_{2,a}/\mathcal{D}_{2,b}$) | 15 / 179 | 32 / 179 | 96 / 179 | 98 / 179 | 151 / 179 | 157 / 179 |
| Sequencer | HiSeq2000/NovaSeq ($\mathcal{D}_{3,a}/\mathcal{D}_{3,b}$) | 1 / 45 | 38 / 45 | 19 / 45 | 29 / 45 | 45 / 45 | 45 / 45 |
| Sequencer | HiSeq2500/NovaSeq ($\mathcal{D}_{4,a}/\mathcal{D}_{4,b}$) | 2 / 45 | 43 / 45 | 40 / 45 | 43 / 45 | 45 / 45 | 45 / 45 |
| Sequencer | HiSeq4000/NovaSeq ($\mathcal{D}_{5,a}/\mathcal{D}_{5,b}$) | 5 / 93 | 86 / 93 | 75 / 93 | 78 / 93 | 93 / 93 | 93 / 93 |
| NovaSeq Chemistry | V1/V1.5 ($\mathcal{D}_{6,a}/\mathcal{D}_{6,b}$) | 12 / 135 | 60 / 135 | 82 / 135 | 89 / 135 | 135 / 135 | 135 / 135 |

TruSeq Nano and Kapa HyperPrep kits. We repeated the experiment done in previous section on each of these groups and reported accuracy in In Table 2. We can observe that our approach drastically improves over standard methods for all groups. In particular, the transport plan Γ inferred by our method perfectly identified the sample pairs for all 6 groups except the TruSeq Nano/Kapa HyperPrep and Kapa/IDT groups, while still improving accuracy by a large margin compared to GC-correction. These results suggest that OT is a suitable framework for estimating patient-to-patient similarities, even in the presence of a limited number of samples (i.e., 45).

### 2.3 Optimal transport disentangles cancer signals from non-biological sources of variation

We further tested the applicability of our method to the detection of haematological cancer and investigated whether data correction preserves the signals of interest (i.e., cancer). For this purpose, we trained simple Machine Learning models using the scikit-learn [52] Python library. The HEMA data set is composed of 179 cases of Hodgkin lymphoma, 22 of multiple myeloma and 37 of diffuse large B-cell lymphoma at different stages, as well as two control sets of size 242 and 257 respectively. The cancer samples as well as the healthy cohort from *D*_7_ have been processed with the TruSeq ChIP kit (Illumina), while the second healthy set from *D*_8_ has been prepared with the TruSeq Nano kit (Illumina). TruSeq Nano samples have been corrected to match the distribution of the TruSeq ChIP controls and validation was performed on the TruSeq ChIP cases and controls. As explained in the Methods section and as illustrated in Fig. 6D, corrected samples were used only for training, to avoid overoptimistic estimation of sensitivity and specificity resulting from controls being accidentally shifted away from the cancer cases.

In Table 3, we reported the performance of binary prediction of haematological cancers, namely Hodgkin lymphoma (HL), diffuse large B-cell lymphoma (DLBCL) and multiple myeloma (MM). Evaluation metrics have been estimated using fivefold cross-validation. Only samples from the training set have been corrected by our method, so as to avoid any data contamination between the training and validation set (see Methods). Sensitivity, specificity and MCC were determined based on the cutoff that produced the highest MCCs. As can be observed, data correction with our domain adaptation approach almost systematically improves cancer detection in terms of MCC, AUROC and AUPR, either through an increase in sensitivity, specificity, or both. In particular, it produced the best MCC in all of the nine settings (3 models *×* 3 pathologies), the best AUPR in 8 settings and the best AUROC in 7 settings. Strikingly, it outperformed GC-correction by 8.6% and 5.3% in MCC for DLBCL prediction with logistic regression and random forest, respectively.

**Table 3:** Haematological cancer detection using supervised approaches. Sensitivity, specificity, Matthews correction coefficient (MCC), AUROC and AUPR obtained through validation of 3 supervised models. These models have been successively trained to distinguish Hodgkin lymphoma, DLBCL and multiple myeloma cases from healthy controls. Sensitivity, specificity and MCC were computed using the cutoff that maximises MCC.

| Supervised model | Logistic regression |  |  |  |  | Random forest |  |  |  |  | Support vector machine |  |  |  |  |
| --- | --- | --- | --- | --- | --- | --- | --- | --- | --- | --- | --- | --- | --- | --- | --- |
| Metric | Sen. | Spec. | MCC | AUROC | AUPR | Sen. | Spec. | MCC | AUROC | AUPR | Sen. | Spec. | MCC | AUROC | AUPR |
| Hodgkin lymphoma |  |  |  |  |  |  |  |  |  |  |  |  |  |  |  |
| No correction | 95.0 % | 97.1 % | 92.2 % | 98.2 % | 98.3 % | 76.0 % | 97.5 % | 76.9 % | 93.3 % | 93.4 % | 95.5 % | 86.0 % | 80.6 % | 96.3 % | 95.4 % |
| Center-and-scale | 96.1 % | 94.6 % | 90.4 % | 98.3 % | 98.3 % | 89.9 % | 89.3 % | 78.8 % | 95.0 % | 94.1 % | 92.7 % | 88.8 % | 80.9 % | 95.1 % | 93.8 % |
| GC correction | 92.7 % | 99.2 % | 92.8 % | 98.7 % | 98.8 % | 87.7 % | 97.5 % | 86.5 % | 97.0 % | 97.1 % | 96.1 % | 95.9 % | 91.8 % | 98.7 % | 98.6 % |
| Optimal transport | 96.6 % | 97.1 % | 93.7 % | 98.9 % | 99.0 % | 92.2 % | 97.1 % | 89.8 % | 97.8 % | 97.9 % | 94.4 % | 97.5 % | 92.2 % | 98.6 % | 98.3 % |
| DLBCL |  |  |  |  |  |  |  |  |  |  |  |  |  |  |  |
| No correction | 75.7 % | 99.6 % | 83.6 % | 92.8 % | 86.1 % | 32.4 % | 99.2 % | 49.1 % | 84.2 % | 56.2 % | 78.4 % | 97.5 % | 77.7 % | 95.5 % | 83.9 % |
| Center-and-scale | 86.5 % | 98.8 % | 87.3 % | 96.6 % | 92.8 % | 64.9 % | 97.5 % | 68.3 % | 94.1 % | 79.0 % | 89.2 % | 95.9 % | 79.9 % | 96.8 % | 84.2 % |
| GC correction | 81.1 % | 98.3 % | 82.3 % | 96.4 % | 87.7 % | 67.6 % | 97.9 % | 71.7 % | 91.5 % | 78.6 % | 78.4 % | 99.2 % | 83.7 % | 97.4 % | 88.0 % |
| Optimal transport | 94.6 % | 98.3 % | 90.9 % | 97.9 % | 94.4 % | 73.0 % | 98.3 % | 77.0 % | 94.6 % | 86.0 % | 89.2 % | 97.5 % | 84.8 % | 96.7 % | 88.1 % |
| Multiple myeloma |  |  |  |  |  |  |  |  |  |  |  |  |  |  |  |
| No correction | 90.9 % | 99.6 % | 92.4 % | 98.0 % | 92.1 % | 63.6 % | 98.8 % | 70.3 % | 91.0 % | 70.6 % | 63.6 % | 96.7 % | 60.3 % | 93.7 % | 62.4 % |
| Center-and-scale | 72.7 % | 99.6 % | 81.4 % | 97.3 % | 89.4 % | 63.6 % | 98.3 % | 68.0 % | 93.3 % | 64.6 % | 59.1 % | 97.5 % | 60.5 % | 93.6 % | 63.1 % |
| GC correction | 90.9 % | 100.0 % | 95.0 % | 99.2 % | 96.7 % | 77.3 % | 100.0 % | 87.0 % | 95.5 % | 89.4 % | 81.8 % | 97.9 % | 78.2 % | 97.4 % | 83.6 % |
| Optimal transport | 95.5 % | 100.0 % | 97.5 % | 99.7 % | 98.0 % | 86.4 % | 99.6 % | 89.8 % | 96.1 % | 91.6 % | 95.5 % | 95.9 % | 78.4 % | 98.3 % | 85.7 % |

Analogous results obtained on the OV data set have been reported in Table 4. Because the ovarian data set contains cases and controls from both domains, we could perform our validation in both directions (adapting samples from *D*_9_ to *D*_10_ and assessing performance on remaining samples from *D*_10_, and vice versa), corresponding to the top and bottom parts of Table 4. The proposed method systematically produced an improvement in MCC and AUPR in all three settings and improved AUROC in two out of the three settings. In particular, we noticed gains of 2.8%, 6.8% and 8.8% in MCC, respectively. In light of these results, we showed the ability of our method to disentangle sources of variation of biological and technical origins. Indeed, the supervised Machine Learning approaches for cancer detection benefited from the improvement in data quality resulting from domain adaptation, yielding better generalization and validation accuracy. Although these AUROC scores are far from being clinically relevant, they must be contextualized. First, we evaluated our predictive models in an artificially difficult setting where newly-collected samples have been processed with a different technology, while wet-lab protocols should be standardised in clinical settings. Second, cancer cohorts include many low-grade and borderline cases, which are most likely chromosomally stable and therefore may be overlooked by our CNA-based approach.

**Table 4:** Ovarian carcinoma detection using supervised approaches. Sensitivity, specificity, Matthews correction coefficient (MCC), AUROC and AUPR obtained through validation of 3 supervised models. These models have been trained to distinguish ovarian carcinoma cases from healthy individuals. Sensitivity, specificity and MCC were computed using the cutoff that maximises MCC.

| Supervised model | Logistic regression |  |  |  |  | Random forest |  |  |  |  | Support vector machine |  |  |  |  |
| --- | --- | --- | --- | --- | --- | --- | --- | --- | --- | --- | --- | --- | --- | --- | --- |
|  | Sen. | Spec. | MCC | AUROC | AUPR | Sen. | Spec. | MCC | AUROC | AUPR | Sen. | Spec. | MCC | AUROC | AUPR |
| $\mathcal{D}_9 \rightarrow \mathcal{D}_{10}$ | | | | | | | | | | | | | | | |
| No correction | 59.9 % | 80.7 % | 33.4 % | 67.2 % | 92.9 % | 50.5 % | **81.3 %** | 26.0 % | 59.6 % | 90.5 % | 57.5 % | **87.9 %** | 34.5 % | 68.4 % | 93.5 % |
| Centering-scaling | 57.3 % | 81.9 % | 32.7 % | 65.7 % | 92.7 % | 61.5 % | 65.2 % | 28.7 % | 58.4 % | 89.8 % | 64.8 % | 75.5 % | 35.1 % | 67.8 % | 93.2 % |
| GC correction | **60.1 %** | 79.4 % | 33.0 % | 68.0 % | 93.2 % | 64.7 % | 72.0 % | 32.1 % | 67.0 % | 92.7 % | 62.1 % | 80.4 % | 33.3 % | 68.8 % | 93.5 % |
| Optimal transport | 58.5 % | **86.6 %** | **35.4 %** | **70.5 %** | **94.0 %** | **75.5 %** | 64.9 % | **38.0 %** | **70.8 %** | **93.7 %** | **71.4 %** | 74.1 % | **40.0 %** | **73.7 %** | **94.5 %** |
| $\mathcal{D}_{10} \rightarrow \mathcal{D}_9$ | | | | | | | | | | | | | | | |
| No correction | 77.0 % | 70.3 % | 43.3 % | 77.5 % | 93.4 % | **78.6 %** | 77.5 % | **51.0 %** | **83.0 %** | **95.5 %** | **75.8 %** | 81.4 % | 50.4 % | **82.5 %** | 95.5 % |
| Centering-scaling | 77.9 % | 70.6 % | 44.5 % | 77.5 % | 93.5 % | 73.1 % | 84.0 % | 49.0 % | 82.1 % | 95.3 % | 72.1 % | 86.8 % | 49.9 % | 81.9 % | 95.4 % |
| GC correction | **82.2 %** | 68.0 % | 47.3 % | 80.3 % | 94.4 % | 67.4 % | 90.6 % | 48.4 % | 80.0 % | 95.0 % | 70.7 % | 86.7 % | 48.4 % | 81.0 % | 95.2 % |
| Optimal transport | 79.4 % | **78.5 %** | **52.0 %** | **83.3 %** | **95.7 %** | 64.9 % | **93.4 %** | 47.0 % | 78.8 % | 94.7 % | 69.9 % | **91.6 %** | **50.7 %** | **82.5 %** | **95.7 %** |

### 2.4 Optimal transport preserves copy number aberrations across domains

As we are fully aware of the overfitting risks associated with our model, we made sure the adapted samples were consistent with the original data by verifying whether the CNAs of each patient were conserved. For this purpose, clonal and subclonal CNAs were called using the `ichorCNA` v0.2.0 R package (details in Suppl. Mat. 2) and we benchmarked our method on the entire OV data set. Indeed, our domain adaptation method provides adjusted profiles that are transparently interpretable and are directly comparable across domains, allowing their comparison in terms of read counts before and after correction.

Let us denote by *D*_9_ and *D*_10_ the wet labs from which the samples originate (see Table 6), respectively [53] and [54]. We first built a panel of controls (reference set) using the 79 controls from domain *D*_9_ and called CNAs in cancer cases from *D*_9_. Then, we built a panel of controls using the 39 controls from *D*_10_ and called CNAs in the same cancer cases from *D*_9_, after adapting them with our proposed approach to match the distribution of cancer cases in *D*_10_. Finally, we quantified the similarity of `ichorCNA` results using different metrics, as shown in Table 5. Because `ichorCNA` performs GC-correction “under the hood”, we did not include GC-correction in the benchmark, as it would produce results highly similar to the baseline. Also, in the case of center-and-scale standardisation we enforced positivity by clipping the corrected read counts, as negative values cannot be handled by `ichorCNA`.

**Table 5:** Quantitative assessment of the consistency of CNA calling. `ichorCNA` results on ovarian carcinoma cases from *D*_9_ using the panel of controls from *D*_9_, compared to the same cases corrected (*D*_9_ *→− D*_10_) by each method and using the panel of controls from *D*_10_. Metrics in the upper part of the table focus on per-bin metrics, namely the copy number in each bin, the presence of a CNA (copy number *̸*= 2) in each bin, the SOV REFINE [55] score and the log-ratios. We used the SOV REFINE segmentation metric to measure the overlap between called CNAs. The metrics in the bottom section of the table are the average absolute errors on different model parameters estimated by `ichorCNA`.

| Metric | No correction | center-and-scale standardisation | OT |
| --- | --- | --- | --- |
| Accuracy (copy number) | 71.3 % | 39.5 % | **73.4 %** |
| Accuracy (presence of CNA) | 76.8 % | 53.2 % | **78.7 %** |
| SOV_REFINE (copy number) | 0.7401 | 0.5218 | **0.7552** |
| Error on log-ratios | 0.0083 | 0.1164 | **0.0070** |
| Error on tumour fraction | 0.0045 | 0.0197 | **0.0042** |
| Error on tumour ploidy | 0.0880 | 0.2293 | **0.0836** |
| Error on tumour cellular prevalence | 0.0888 | 0.1209 | **0.0866** |
| Error on proportion of subclonal CNAs | 0.1366 | 0.1414 | **0.1268** |

Even in the absence of any correction, the per-bin copy numbers estimated for cancer CNAs are not consistent, as we see that the accuracy and SOV REFINE measures are far from being perfect. This can be attributed not only to (1) the difference in protocols used to produce the two panels of normals, but also (2) the limited number of controls, (3) the fact that the controls differ between the two domains, and (4) the experimental uncertainty (stochastic noise). Overall, both center-and-scale standardisation and optimal transport preserved the original normalised read counts sufficiently since they both provided results similar to the baseline (“no correction”). However, our method improves over center-and-scale standardisation regardless of the evaluation metric. After applying our DA method on the cancer cases, 73.4% of the bins were assigned the correct copy number and segmentation of CNAs produced an overlap score of 0.7552. As illustrated in Fig. 2, the copy numbers called by `ichorCNA` in *D*_9_ (panel A) are mostly preserved without (B) or after (D) correction. The least consistent results were produced by center-and-scale standardisation, where some disruptions have been introduced at multiple locations (e.g., some higher copy numbers in chromosome 3).

**Figure 2:**
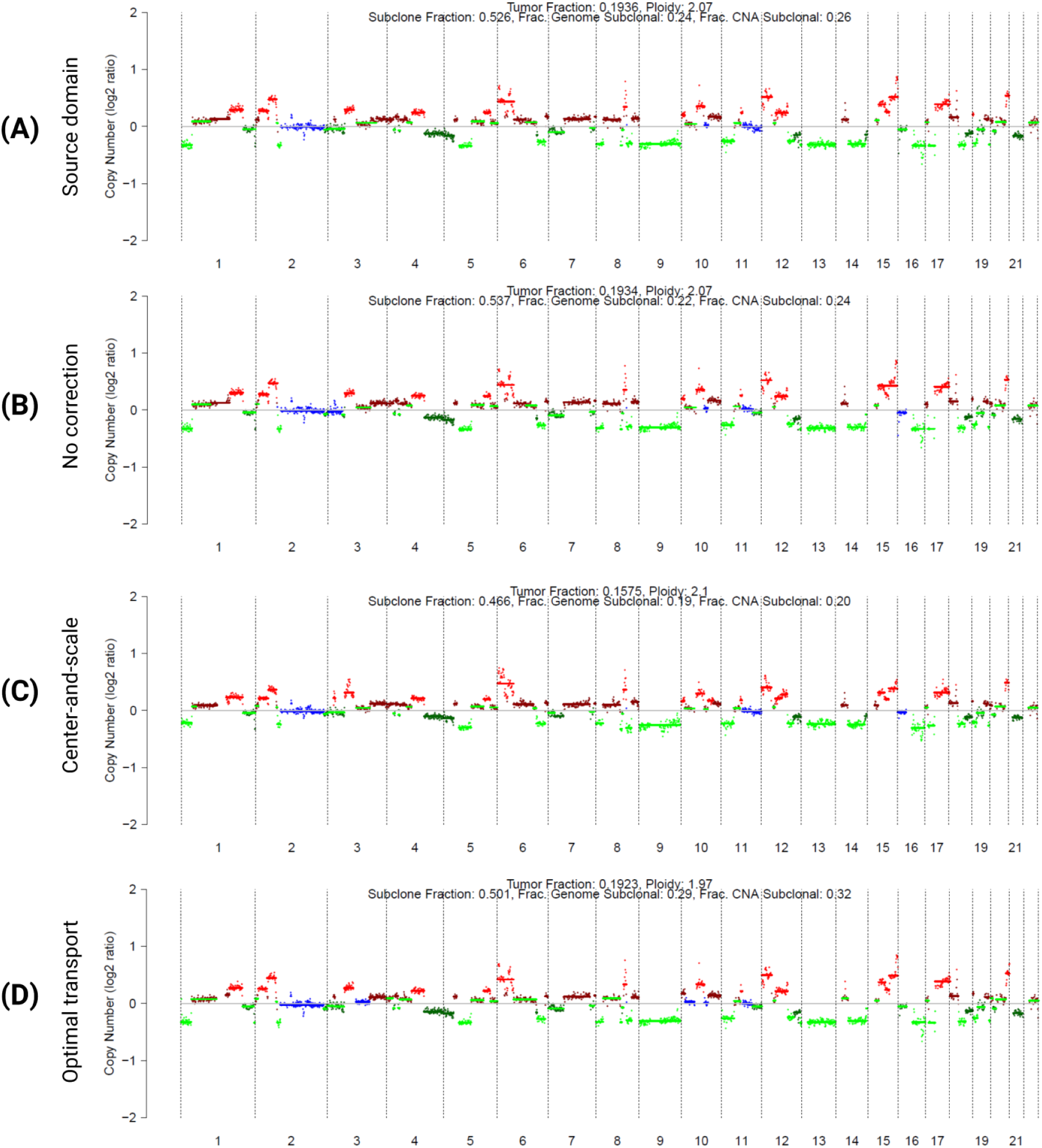
Qualitative assessment of the consistency of CNA calling. Comparison of the CNAs called by `ichorCNA` on a late stage ovarian carcinoma case from *D*_9_, before and after domain adaptation. Green and red colouring correspond to deletions and gains, respectively. (A) Using *D*_9_ controls. (B) Using *D*_10_ controls. (C) *D*_9_ cancer cases (including the case shown) centered-and-scaled toward *D*_10_ and analysed with *D*_10_ controls. (D) *D*_9_ cancer cases OT-corrected toward *D*_10_, analysed with *D*_10_ controls.

The average absolute error on the estimation of tumour fraction is 0.0042, which is acceptable given the error of 0.0045 in the absence of any correction and the standard deviation of the tumour fraction estimates (0.0464). Despite the limited size of our reference sets and therefore the potentially overpessimistic assessment of the inconsistencies of `ichorCNA`’s results, we conclude that most of the CNAs have been preserved and that the proposed method does not disrupt the original data, as a more straightforward standardisation approach would. Since `ichorCNA` offers the possibility to call CNAs without panels of normals, we ran similar analysis without controls and observed more consistent results between the two domains. We reported these results in Suppl. Tab. 1 and Suppl. Fig. 1.

Estimated tumour fractions before and after correction have been reported in Fig. 3, showing good consistency both in the presence (r=0.973, *p*-value=1.09e-205, two-sided test) or the absence (r=0.980, *p*-value=1.77e-224) of the *D*_10_ panel of controls.

**Figure 3:**
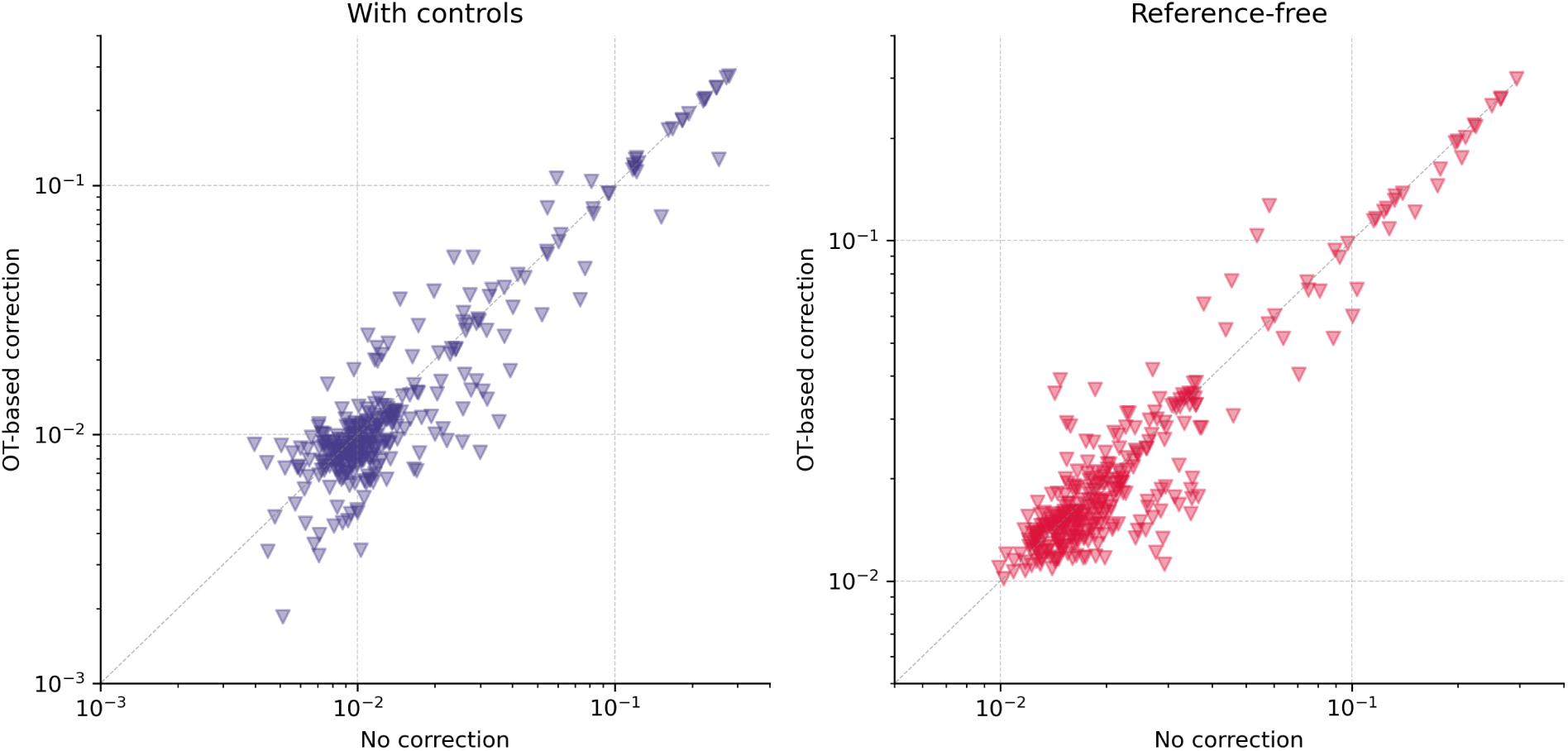
Tumour fractions before and after domain adaptation. Fractions have been estimated with `ichorCNA` before and after adapting the *D*_9_ ovarian carcinoma cases toward the *D*_10_ cases. Results have been produced both with (Left) and without (Right) the panel of controls from domain *D*_10_. Fractions are shown in log-scale.

### 2.5 Preanalytical variables are mostly discrete

A major limitation of our approach is its inherent restriction to discrete settings, where the technical counfounder is acting as a dummy variable and reflects whether some technology as been used to produce a sample or not. However, to the best of our knowledge there is no continuous preanalytical variable in whole-genome sequencing that induces gradual changes in the normalised read counts and in our data sets. A potential exception is the plasma separation delay, measured as the time elapsing between the blood draw and the separation of the plasma from the buffy coat. We tested Pearson and Spearman correlation for each 1 Mb genome bin on the HEMA data set, using a significance level of 0.01 and applying the Benjamini-Hochberg procedure to account for multiple testing. As shown in Fig. 4B, no bin was found to be significantly correlated with the plasma separation delay. The normalised reads counts of the NIPT samples in the first 1 Mb bin of chromosome 6 (bin showing highest correlation with plasma separation delay) are shown in fig. 4A.

**Figure 4:**
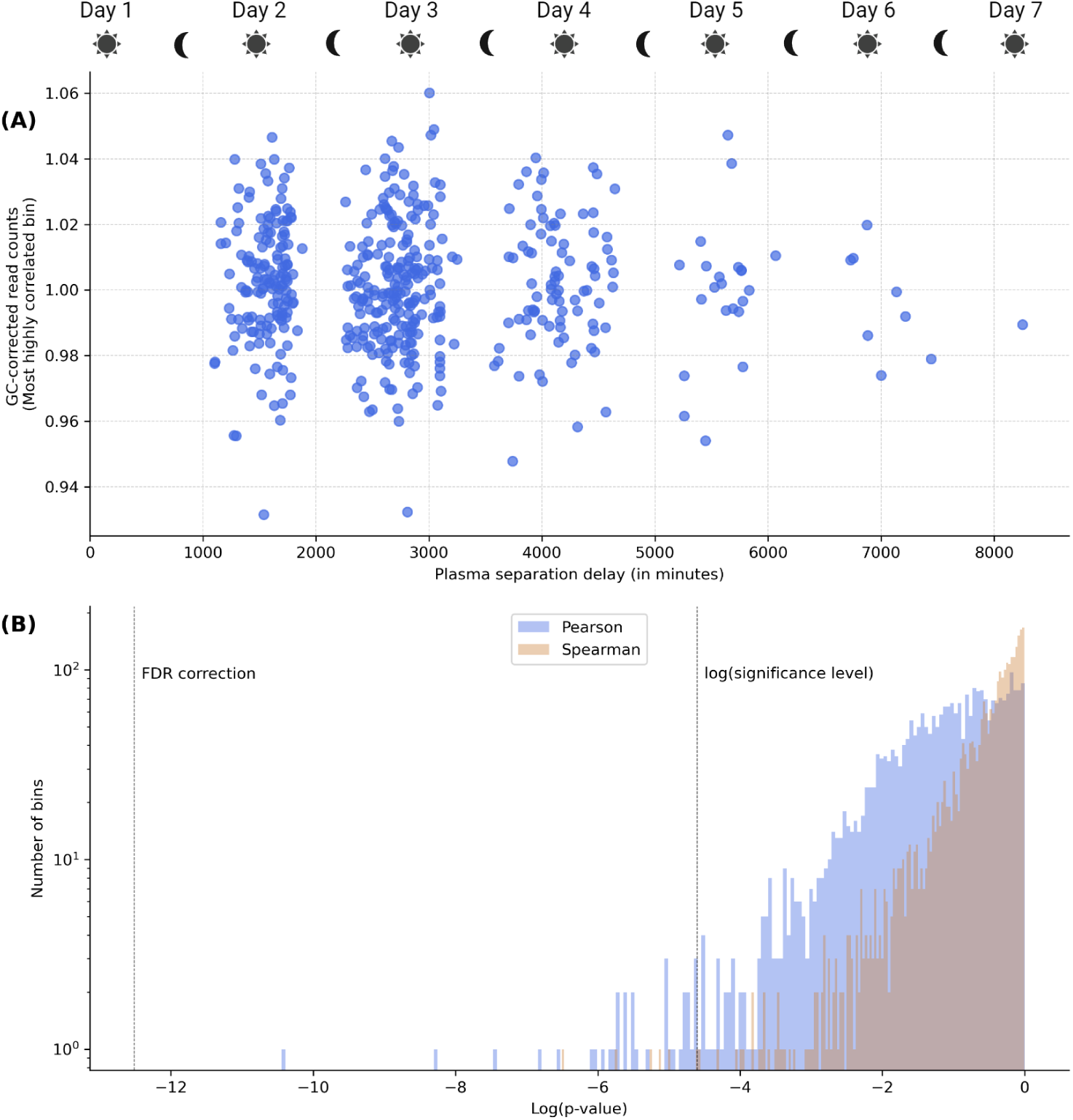
Effect of plasma separation delay on coverage profiles. (A) GC-corrected normalised read counts of all samples from the NIPT data set for a specific bin (first 1 Mb bin from chromosome 6), namely the one giving the strongest linear correlation with the plasma separation delay (Pearson’s *r*=0.1838, *p*-value=1.38e-5, two-sided test). (B) Distribution of the *p*-values computed likewise for each 1 Mb bin, and reported as a histogram. Histogram is shown in log-scale.

### 2.6 GC-correction is not sufficient to decorrelate read counts from GC-content

In Figure 1, we showed that GC-correction did not help in reducing the dissimilarity between the two control sets from the HEMA data set. We also reported similar results on the two other data sets in Suppl. Mat. 3. While GC-correction succeeds at reducing the individual variability (experimental variance) of samples as shown by the improved accuracy in Tables 1 and 2, it fails at alleviating the biases introduced by changes in sequencer or library preparation method. Indeed, while this approach improves cancer detection on average by removing technical variations based on GC-content, it does not systematically produces performance gains, does not efficiently capture similarities between profiles originating from the same biological sample and does not completely remove the clustering effects introduced by the changes in the sequencing methodology. By contrast, our method showed that these expectations can be met through the modelling of patient-to-patient similarities and explicit constraining of the samples based on quantiles. Indeed, these latter constraints drastically lower the risks of overfitting and ensure that the mapping between the cohorts is performed in a biologically meaningful manner.

In Figure 5, we reported the two-sample Kolmogorov-Smirnov *p*-values from Figure 1b as a function of the GC-content, as well as the median *p*-value per 0.5% GC stratum. Normalised read counts (top left panel) exhibited strong relationship between median *p*-values and GC-content, demonstrating that regions with low and high GC-content are the most biased by the change from the KAPA HyperPrep to the

**Figure 5:**
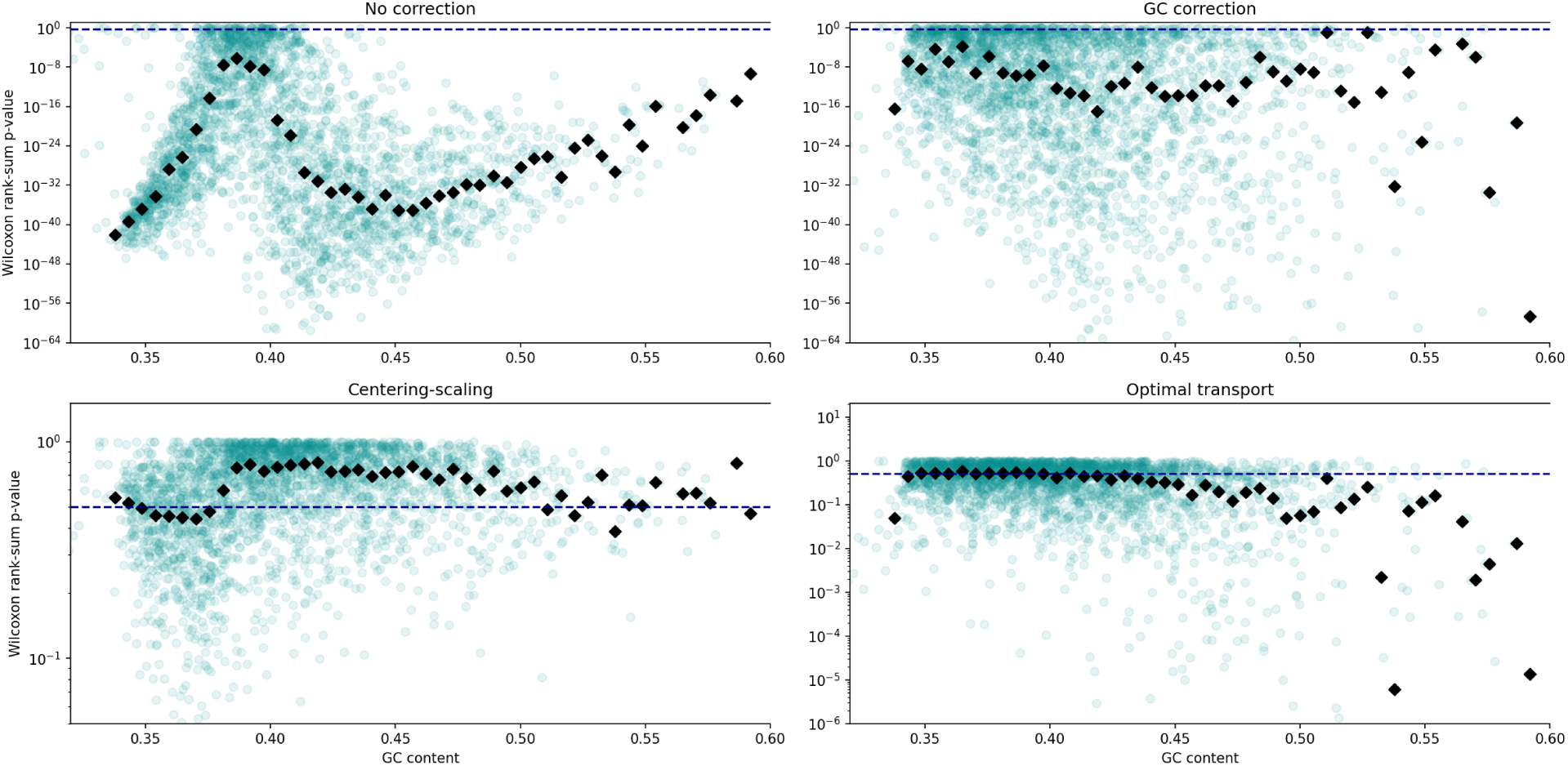
Coverage difference between domains as a function of GC-content. Two-sample Kolmogorov-Smirnov testing, for each 1 Mb bin, of the difference between the two control sets from the HEMA data set processed with different library preparation kits. *p*-values are shown in log-scale, as a function of the GC-content of each bin. Dashed line corresponds to a 0.5 *p*-value and black markers to the median *p*-values per 0.5% GC stratum.

TruSeq ChIP library preparation kit. While center-and-scale standardisation is capable of centering these *p*-values around 0.5 (which is expected under the null hypothesis since *p*-values are uniformly distributed in the [0, 1] interval), we observed the same trend. GC-correction drastically improved in that respect, as no clear correlation can be observed. However, the median *p*-value still varies from stratum to stratum, suggesting that some subtle and nonlinear GC biases remained. Finally, our proposed approach produced the most consistent results across the GC-content values, showing that it more effectively alleviated these residual biases.

### 2.7 Domain expertise is the best regularisation

There are multiple mechanisms put in place within our model to constrain it to infer a matrix *X* that is as meaningful as possible, namely the quantile-based regularisation function, the positivity constraint on the read counts, the median normalisation and GC correction. While these constraints are not guarantees of performance *per se*, they restrict the size of the search space drastically, eliminating a large proportion of irrelevant solutions. While our method was originally designed for correcting normalised read counts with the detection of CNAs in mind, it remains sufficiently generic to be applied to any (whole-)genome sequencing or array-based data set. Indeed, the only requirement is the representativeness of the cohorts in all domains. However, the requirements imposed on the data are problem-dependent and heavily depend on the nature of the data sets. Therefore, from a general perspective, extra care should be given to the assumptions underlying the model. For instance, preserving the quantiles (e.g., *z*-scores) might not necessarily be desirable, as read counts are heavily subject to noisy fluctuations that should preferably not be transferred from domain to domain.

Beyond aforementioned limitations, our findings open new perspectives for the analysis of highdimensional whole-genome sequencing data and suggest that appropriate modelling of technical confounders enables the joint analysis of cohorts sequenced at different points in time (changes of sequencing platform, library preparation kit, DNA extraction method) and space (team, hospital, country). Finally, the analysis of larger data sets is expected to strengthen the detection power of statistical models based on cfDNA data and enable the presymptomatic detection of more subtle cancer signals.

## 3 Methods

Throughout the paper, we referred to *cohort* as a set of samples sharing similar high-level characteristics (e.g., a set of healthy controls, a set of pregnant women, a set of ovarian cancer patients) *and* processed using similar protocols. A *domain* is a set that can include multiple cohorts, with no regard for the biological state as only the protocol is considered. Finally, a *data set* can itself include multiple domains, as each data set has been used to assess our method’s ability to correct for biases between the domains contained in this data set.

### 3.1 Clinical data

We benchmarked our method on three data sets produced in-house, each used for a different purpose. The peculiarities of each data set have been summarised in Table 6. All data sets have a median sequencing coverage between 0.1x and 0.2x.

**Table 6:**
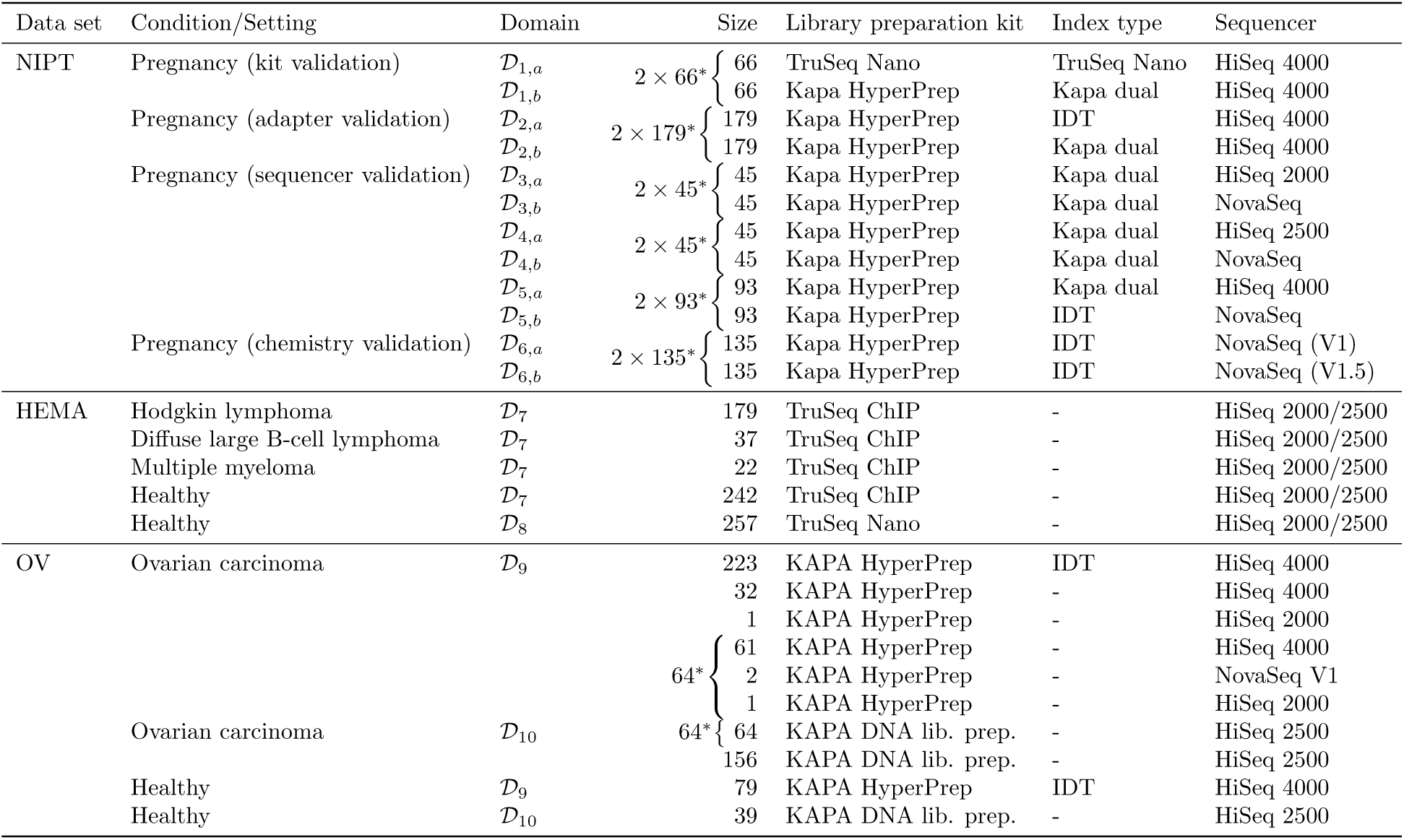
Summary of the data sets used in this study. Samples in sets marked with a ‘*∗*’ have been processed twice, allowing quantitative assessment of the different biases caused by the changes of sequencing protocols. Domains have been defined based on our experiments, as well as the protocol differences shown in the table. For clarity purposes, we systematically refer to these domains in the results section.

The data sets used in the present work have been collected during studies previously approved by the ethical committee of the University Hospitals Leuven under S/57999, S/62285, S/62795, S/50623, S/56534, S/63240, S/51375, S/55904, S/57144, S/59207, S/64205 and S/64035. Blood samples were collected either into Streck cfDNA BCT or Roche Cell-Free DNA Collection Tubes. cfDNA was extracted using either the QIAamp Circulating Nucleic Acid Kit or the Maxwell automated protocol. Samples were pooled by batches of 20 for multiplex sequencing using all lanes of Illumina flow cells. Each pool was sequenced either on the Illumina HiSeq 2000, HiSeq 2500, HiSeq 4000 or NovaSeq 6000 platform, single-end 1×36bp, 1×50bp or paired-end 2×50bp.

The first data set consists of 563 validation samples collected in the context of Non-Invasive Prenatal Testing (NIPT) [56] and processed within the standard NIPT routine twice. These paired samples are divided in 6 validation groups (2 *×* 6 domains), each used to quantify the distributional shift introduced by the change of *one* preanalytical variable. The libraries of 66 biological samples have been prepared with either the TruSeq Nano DNA Sample Preparation Kit (Illumina) or the KAPA HyperPrep Kit (Roche) with Kapa Dual indexed adapters. 179 samples have been used with either IDT indexes or KAPA Dual indexed adapters. 45 samples have been processed either by the HiSeq 2000 or NovaSeq platform. 45 samples have been processed either by the HiSeq 2500 or NovaSeq platform. 93 samples have been processed either by the HiSeq 4000 or NovaSeq platform. Finally, 135 samples have been processed by a NovaSeq platform, with either V1 or V1.5 chemistry. In total, this results in 2 *×* 563 paired samples. We refer to this first data set as NIPT for short.

Our second data set (HEMA) focuses on haematological malignancies and is composed of 179 cases of Hodgkin lymphoma (HL), 37 of diffuse large B-cell lymphoma (DLBCL) and 22 of multiple myeloma, as well as 498 controls. Among those, 177 HL cases and 260 controls have been published in a previous study [57] and the entirety of the haematological cancer cases have been included in one of our studies (GipXplore [53]). The libraries of 242 out of the 499 controls have been prepared with the same kit as the haematological cancer cases, namely the TruSeq ChIP Library Preparation Kit (Illumina) [4]. The remaining ones have been prepared with the TruSeq Nano kit.

Finally, we further validated our OT-based bias removal approach with controls and ovarian carcinoma cases sequenced by two different teams [54] including ours. These samples were not derived from cancer patients with overt clinical disease, but rather the presence of a suspicious malignancy based on imaging. We refer to this last data set simply as OV. Protocols vary in multiple ways. As an example, all of the samples in *D*_9_ (see Table 6) have been processed with HiSeq 2500, while all samples in domain *D*_10_ have been sequenced by an instrument that differed from HiSeq 2500. Samples from *D*_9_ and *D*_10_ have been prepared with the KAPA HyperPrep and KAPA DNA library preparation kits, respectively. Ovarian carcinoma samples from *D*_10_ have been manually extracted with the QIAamp CAN kit. Let’s note that *D*_9_ does not strictly stick to our definition of domain. Despite the heterogeneity caused by the presence of multiple sequencers, we artificially grouped the samples in order to simplify the comparison between laboratories but also better reflect the heterogeneity expected to be encountered in Big Data settings.

### 3.2 Data preprocessing

Reads were first aligned to the reference genome hg38 using the Burrows-Wheeler aligner [58], only considering the 22 autosomes. Then, read duplicates were removed with Picard tools [59] and remaining ones were recalibrated with the Genome Analysis Toolkit [60]. Finally, reads were counted in predefined bins of size 1 Mb. Such size offers a good tradeoff between noise reduction and the granularity of achievable chromosomal aberration detection. Finally, counts were normalised by dividing by the median count per Mb of the whole profile to correct for the effective sequencing depth.

### 3.3 GC-content bias correction

We performed GC-correction by dividing normalised read counts by their estimate according to a Locally Weighted Scatterplot Smoothing (LOWESS) model [61], where the exogenous variable is the GC-content of the bin, and the endogenous variable is the corresponding normalised read count. 30% of the data points (bins) have been used to predict the endogenous variable. We used the Python package statsmodels v0.12.2 [62] to implement the LOWESS correction. We also designed a differential version of GC-correction that is PyTorch-compliant and used by our DA method to ensure that the reads counts of adapted cohorts do not correlate with GC-content. More details are provided in Suppl. Mat. 1.

### 3.4 Center-and-scale standardisation

We benchmarked our method against a more straightforward approach, consisting in the standardisation of each cohort or data set separately. This approach ensures that the *z*-scores are centered in all domains and allows comparability as long as each cohort is representative of the population. For each 1Mb bin, we centered and scaled the normalised read counts as follows:

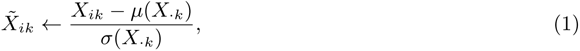

where *µ*(*X_·k_*) is the median of normalised read counts within bin *k* across all samples from the same cohort and *σ*(*X_·k_*) the square root of the average squared deviation from this median. The median has been used in place of the mean for robustness against outliers, such as profiles with aberrant CNAs. We refer to this method as center-and-scale standardisation throughout the manuscript.

### 3.5 Domain adaptation using optimal transport

We defined the best correction function as the one that minimises statistical dissimilarity metric between two cohorts. Given the multivariate nature of the problem and the strong mathematical foundations behind optimal transport, we propose to use the Wasserstein distance to quantify the discrepancy between data sets.

The general principle of OT is to match two probability distributions by transporting the probability mass of one distribution into the other with minimal effort (hence the name *optimal transport*). Distributions that can be transported into each other at a low cost are considered highly similar. In the case of discrete samples, OT amounts to finding a discrete probabilistic mapping (called the *transport plan*) of the source samples onto the target samples where the mapping of a source sample to a target sample bears some associated cost. We consider thus two data matrices 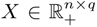 and 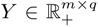, as illustrated by the normalised read counts matrices in Fig. 6A, where *n* and *m* are the sample sizes of the each domain and *q* is the number of predefined bins. As samples are all assumed to be of equal importance, we choose uniform probabilistic weights *νi* = 1*/n, ∀i* and *µj* = 1*/m, ∀j* to define a probability distribution on these discrete samples (assuming no two samples can be identical). We also consider the associated pairwise Euclidean distance matrix *C ∈* R*^n×m^*. For normalised read counts, the matrix *Y* is first GC-corrected. The matrix *X* will be GC-corrected during OT, as part of a joint optimisation process. The reason for not correcting *X* upfront is that OT might artificially introduce correlations with GC-content *a posteriori*. Instead, we implemented median-normalisation and GC-correction as a differentiable function *f* (see Suppl. Mat. 1).

The Wasserstein distance is defined, in its discrete form, by

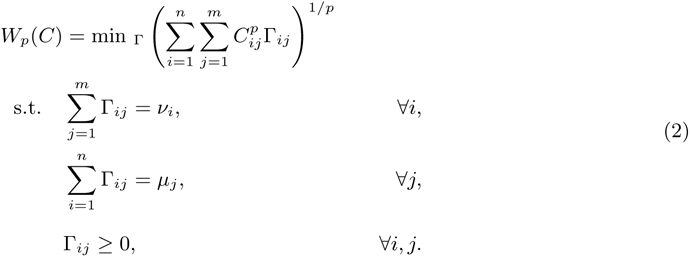

Matrix Γ, usually referred to as the transport plan and depicted in Fig. 6B, can be interpreted as the amount of probability mass transferred from points of the source domain to the target domain through optimal transport. In particular, Γ*ij* is the probability mass transferred from point *i* in the source domain to point *j* in the target domain. Such interpretation allows us to use Γ as a pairwise similarity matrix and express the domain adaptation problem as a multivariate regression problem. By choosing *p* = 2 and the Euclidean metric as function *d*, as well as by accounting for the reduction of variance caused by *Gamma*, and attaching equal importance to the bins (see details in Suppl. Mat. 1.2), the optimisation problem becomes

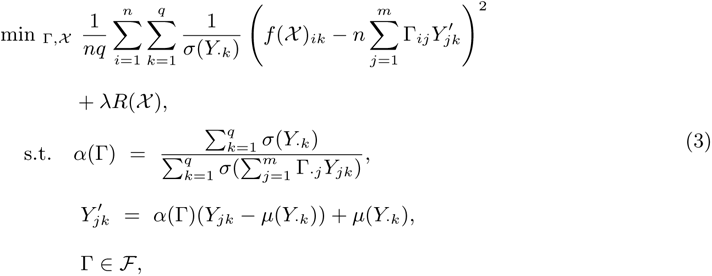

where *α*(Γ) is the variance correction factor, *R*(*X*) is a regularisation term, and *λ* is the associated regularisation hyper-parameter.

The output of our algorithm is matrix *X*, which we interpret as the surrogate of *X* in the target domain.

#### 3.5.1 Regularisation function

While the Wasserstein distance is often supplemented with a regularisation term based on the entropy of Γ [63], we noticed that entropic regularisation tends to reduce the variance of the adapted samples, ultimately collapsing them onto their centroid. This is not a desirable property because in actual high-dimensional data the curse of dimensionality will naturally keep data points distant from each other in the presence of noise. This creates an obstacle to the idea of mapping a source sample to the “closest” target samples. Therefore, we do not regularise the Wasserstein distance based on entropy. Instead, we propose a more informative approach where the deviations (e.g., chromosome gains or deletions) of samples *X* from some reference should be preserved throughout the whole adaptation process.

The regularisation function is defined as a mean squared error function:

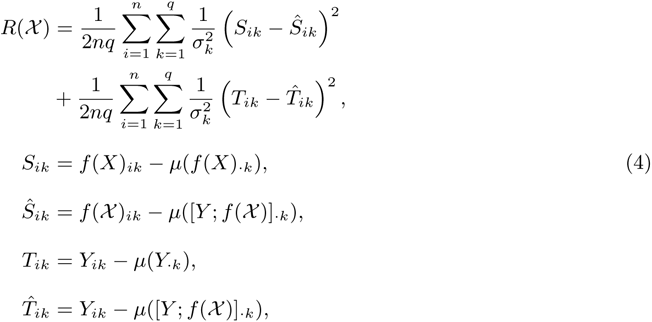

where *f* (*X*) is our differentiable GC-correction function, applied independently on each row of *X*. For a more robust estimation, we computed *µ*(*X_·k_*) as the median over bin *k* (rather than the mean). We used the MATLAB notation [*Y*; *f* (*X*)] to denote the vertical concatenation of matrices *Y* and *f* (*X*) (resulting in a matrix of dimension (*n* + *m*) *× q*). This regularisation function is meant to preserve the quantiles (akin to the *z*-scores) across the two domains. In particular, the first term enforces the consistency of the samples from the source domain, while the second term operates similarly on the samples from the target domain.

#### 3.5.2 Overfitting and stopping criterion

Since our inference process is iterative, a convergence criterion is required in order to end it and limit overcorrection risks. Since our goal is to merge cohorts in such a way that they appear to be drawn from the same distribution, we performed per-bin statistical tests. At each iteration, a *p*-value is computed based on the two-sample Kolmogorov-Smirnov test for each 1 Mb genome bin. Because *p*-values are randomly and possibly uniformly distributed under the null hypothesis [51] (that the two cohorts are representative of the same population), we assumed that the median of these *p*-values should be close to 0.5. In practice, we interrupt the optimisation process as soon as the median *p*-value exceeds this cutoff.

The convergence and its speed are to a large extent impacted by the regularisation rate *λ*, which needs to be picked carefully. When picking *p* = 1 and choosing the squared Euclidean distance as function *d*, then *Wp*(*C*) and *R*(*X*) are both average squared error functions and expected to have similar orders of magnitude after convergence. For this reason, a reasonable *a priori* choice for *λ* would be 1. However, we observed that this default value results in many situations where the model never satisfies the convergence criterion and ends up with a low median *p*-value (e.g., 0.25), despite a large number of iterations (*>* 1000). In many other situations, the model rapidly converges but introduces disruptions in the data, resulting in drastic information loss. Therefore, we chose to make *λ* adaptive and lower it every *e* iterations until the convergence criterion is met. Our motivation is to use the largest value for *λ* (to preserve the original data to the best extent possible) while ensuring that the two data distributions are no longer distinguishable. We arbitrarily chose an initial value *λ*0 = 1000 and reduced *λ* by half every *h* = 20 iterations. Minimal value of *λ* was set to 1 in order to prevent overfitting risks in case where the model fails at reaching the convergence threshold.

### 3.6 Model validation and performance assessment

#### 3.6.1 Downstream supervised learning

Domain adaptation aims at superimposing the data distributions originating from different domains. While this superimposition can be quantified through clustering metrics or visually assessed using kernel principal component analysis or *t*-SNE for example, additional validation is required to ensure that the predictive signal for the malignancies of interest has not been removed during the adaptation process. For this purpose, we trained widely used Machine Learning models for the detection of these malignancies before and after adaptation, using default hyperparameters. We trained logistic regressions, random forests and kernel support vector machines using the scikit-learn [52] Python package.

However, regular validation approaches, such as *k*-fold or leave-one-out cross-validation do not suit our setting, as they may show overoptimistic performance because of contamination between the training and the validation set. Indeed, the domain adaptation model should not be exposed to the validation set, since in the presence of overfitting some corrected samples from the training set will resemble samples from the validation set. Therefore, we propose a problem-specific validation method that excludes from the validation set any sample that does not belong to the target domain, as well as any sample that has been seen by the DA algorithm. During validation, adapted samples are only used to train the model and are not allowed to be left out, therefore not contributing to the estimation of performance. The aim is to prevent the supervised models to correctly assign a label (healthy/cancer) to the left-out sample just because the latter has been shifted arbitrarily far away from the real data distribution by the domain adaptation model. Such procedure can be repeated multiple times on random subsets, similarly to *k*-fold and leave-one-out cross-validation. The subsets generated during our validation procedure are illustrated in Fig. 6D.

**Figure 6:**
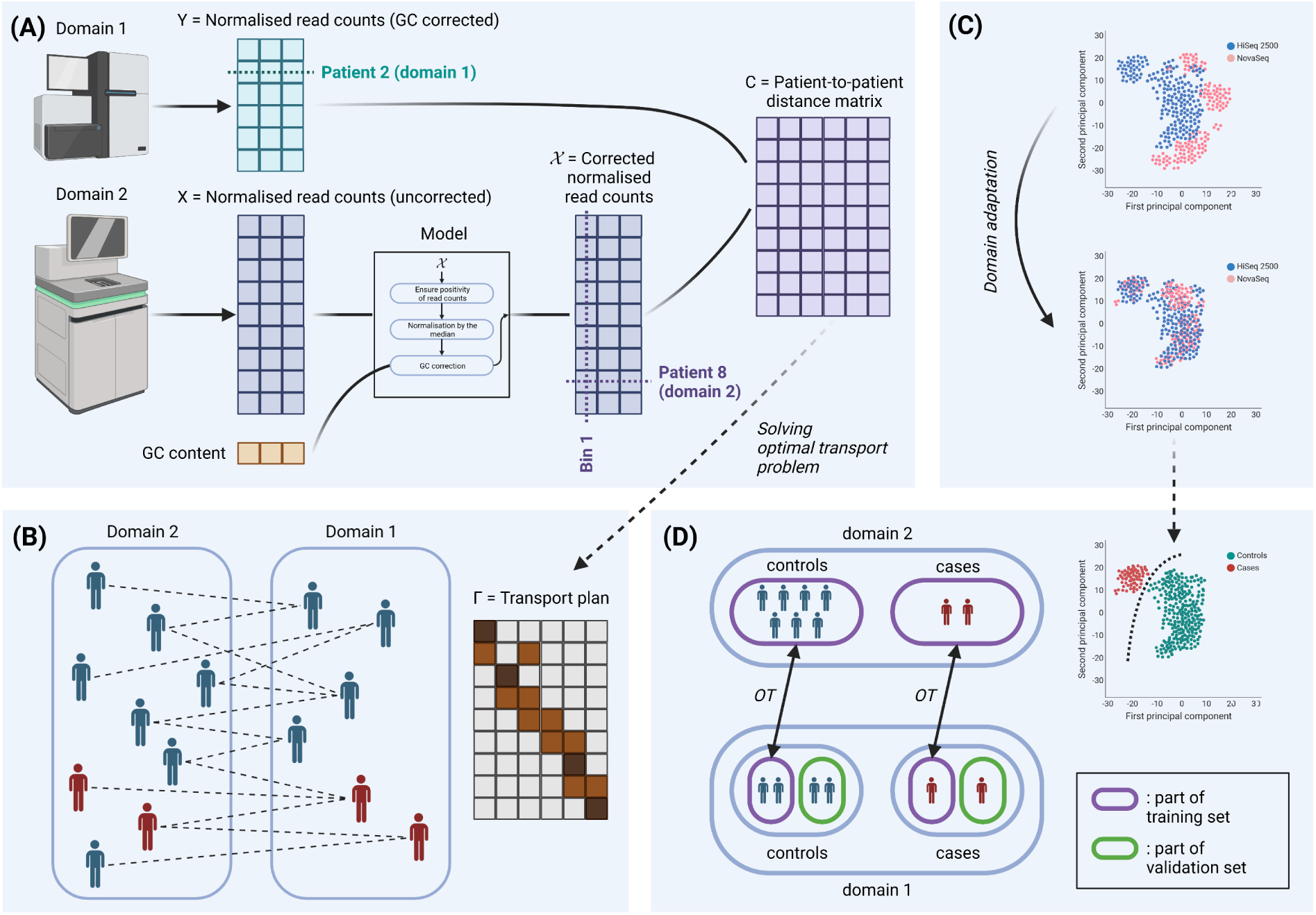
Illustrative summary of our methods. (A) Given two cohorts of cfDNA samples differing by the sequencing pipeline that processed them, the model corrects the second cohort to match the distribution of the first one. After correction, the cost matrix for our OT problem is given by the pairwise Euclidean distances. (B) The solution of the OT problem, named transport plan, assigns patients from Domain 2 to similar patients in Domain 1. The model parameters are found by minimising the Wasserstein distance, as defined by the cost matrix and transport plan. (C) After inference, the two cohorts are merged and ready for downstream analysis. (D) Depiction of the validation procedure used for the purpose of this study.

Since the HEMA data set contains controls and cases in both domains, the validation has been performed in both directions: mapping samples from first domain to the second one before validating in the second one, and vice versa.

Performance of the supervised models for cancer detection was quantified using widely used metrics such as sensitivity, specificity, Area Under the Receiver Operating Characteristic curve (AUROC), Area Under the Precision-Recall curve (AUPR) and the Matthews Correlation Coefficient (MCC).

#### 3.6.2 Evaluation of sample-to-sample mapping using paired samples

For the cohorts in which biological samples have been sequenced twice, we applied OT and assessed whether the distance between paired samples was indeed lowered by the adaptation process. More specifically we computed accuracy, measured as the percentage of profiles from the target domain correctly assigned to the corresponding profile in the source domain, using the closest profile (according to the Euclidean metric) as predictor. By construction, random counterpart assignment would result in a 1*/n* accuracy.

#### 3.6.3 Downstream copy number aberration analysis

Finally, we assessed the ability of the different methods to preserve the copy number aberrations present in the original data. For this purpose, we built a reference set made of the 79 controls from domain *D*_9_ (OV data set, see Table 6) and called CNAs in the cancer cohort from *D*_9_, same data set. Next, we built a reference set based on the 39 controls from *D*_10_ and called CNAs in the same cancer cohort from *D*_9_, after applying our tool to adapt them towards *D*_10_. CNA calling was performed with `ichorCNA`, which would be theoretically expected to produce similar results in the two settings. Therefore, we quantified the agreement between `ichorCNA` results using accuracy and SOV REFINE score [55] between the estimated per-bin copy numbers. Also, we computed the average absolute deviations in the parameters inferred by `ichorCNA`, including tumour fraction or tumour cellular prevalence. Since the tool also proposes to call CNAs in a reference-free fashion, we also conducted the same analysis without the two panels of normals.

## Data availability

Haematological cancer cases and healthy controls constituting the HEMA data set are available from ArrayExpress (https://www.ebi.ac.uk/arrayexpress/) under accession number E-MTAB-10934 as part of the GIPXplore study.

Ovarian carcinoma and healthy controls (OV data set) from domain *D*_10_ of have been previously deposited at the European Genome-phenome Archive (EGA) under study no. EGAS00001005361.

The remaining samples (domain *D*_9_ + NIPT dataset) are in-house cohorts.

Coverage profiles for all the samples have been compiled and uploaded to FigShare (DOI: 10.6084/m9.figshare.24459304).

## Code availability

Our tool is available as an open source package at https://github.com/AntoinePassemiers/DAGIP.

## Acknowledgments

Antoine Passemiers is funded by a Research Foundation – Flanders (FWO) doctoral fellowship (1SB2721N). Tatjana Jatsenko is funded by Agentschap Innoveren en Ondernemen (VLAIO; Flanders Innovation & Entreprenership grant HBC.2018.2108). Joris Robert Vermeesch is funded by FWO (G080217N, S003422N) and KU Leuven (no. C1/018). Daniele Raimondi is funded by a FWO post-doctoral fellowship (12Y5623N). Yves Moreau is funded by (1) Research Council KU Leuven: Symbiosis 4 (C14/22/125); CELSA Active Learning, (2) Innovative Medicines Initiative: MELLODY, (3) Flemish Government (ELIXIR Belgium, IWT, FWO 06260, FWO SBO MICADO S003422N, VLAIO ATHENA HBC.2019.2528) and (4) Impulsfonds AI: VR 2019 2203 DOC.0318/1QUATER Kenniscentrum Data en Maatschappij. An Coosemans is funded by the Flemish Cancer Society (2016/10728/2603). ctDNA samples within the Trans-IOTA study were financed by Kom Op Tegen Kanker (Stand Up to Cancer).

The resources and services used in this work were provided by the VSC (Flemish Supercomputer Center), funded by the Research Foundation - Flanders (FWO) and the Flemish Government. Figure 6 has been created with BioRender.com.

## Author contributions

Designed experiments: A.P, T.J, J.R.V, P.B, A.C, D.T, D.L. Designed computational methods: A.P, Y.M, D.R. Data collection and analysis: A.P, A.V, T.J. Performed computational experiments: A.P. Wrote the first draft of the manuscript: A.P, T.J, D.R, Y.M, J.R.V Revised and approved manuscript: all authors.

